# Degradations of tannin and saponin during co-composting of shell and seed cake of Camellia oleifera Abel

**DOI:** 10.1101/652560

**Authors:** Jinping Zhang, Yue Ying, Xiaohua Yao

**Affiliations:** Research Institute of Subtropical Forestry Chinese Academy of Forestry

**Keywords:** *Camellia oleifera* Abel, shell, seed cake, compost, tannin, saponin, degradation

## Abstract

The degradation processes were studied in this paper of tannin and saponin during the co-composting of the shell and seed cake of *Camellia oleifera* Abel. Four treatments were designed with the dry weight of the seed cake accounting for 1/3(A1), 1/4(A2), 1/5(A3), and 1/10(A4) of the shell. The proportion of the seed cake is positively correlated with the duration of thermophilic phase, highest temperature and degradation rate of tannin and saponin whose maximum were in A1, but negatively correlated with C/N ratio and tannin content which were least in A1 of the final products. The content of saponin were all about 2% finally. The final content of saponin and tannin decreased 68.92-75.22% and 34.57-59.52%. The organic matters, total nutrient (N, P_2_O_5_ and K_2_O) increased with the rising proportion of the seed cake. Overall, the addition of the seed cake promoted the stability, fertilizer efficiency and safety of the co-compost product.

## Introduction

*Camellia oleifera* Abel (*C. oleifera*) is known as one of the top four woody edible oil tree species in the world, and a major economic crops in southern China [1]. By the end of 2017, the cultivation area of *C. oleifera* has expanded to 4.37 million hectares, with annual production of *C. oleifera* seed, oil, shell and seed cake reaching to 2.4, 0.6, 3.6 and 1.8 million tons, respectively, in China. *C. oleifera* shell accounts for more than 60% of the weight of the fresh fruit [2]. In addition to lignocellulose that accounts for 80% of the composition, *C. oleifera* shell also contains saponin and tannin [3] that can be degraded very slowly under natural conditions. *C. oleifera* seed cake is the residue left from the oil extraction, which is mainly composed of crude protein, crude fat, crude fiber, saponin, tannin, ash, caffeine, etc. [4]. The direct disposal or utilization of the *C. oleifera* seed cake can cause serious environmental pollution.

The contents of tannin and saponin in *C. oleifera* shell are about 2.26% and 4.8%, respectively [3], and the saponin content in *C. oleifera* seed cake is about 15-25% [5]. Tannin is a class of defensive secondary metabolites [6] that are produced during the long-term evolution of plants to prevent infection by animal feeding and pathogenic microorganisms, which can inhibit microbial growth and biodegradation [7]. The tannins in *C. oleifera* shell and seed cake are highly polymerized natural polyphenols that can bind to proteins or enzymes, impede the metabolism of microorganisms and inhibit the growth of microorganisms. However, some microorganisms are resistant to tannins, and even can utilize tannins as the sole carbon source [8,9]. Tea saponin is a class of pentacyclic triterpenoids with a β-fragrant tree skeleton. They can be considered as a derivative of oleanane with a polyhydroquinone pentacyclic ring [10]. Tea saponin is highly toxic for cold-blooded animals [11] and can hemolyze red blood cells [12]. The safe intake for warm-blooded animals is 50-150 mg/kg per day, and the effective concentration for killing *lumbricina* is 0.3 mg/ml. Tea saponin can function as the antifeedant of Plutella xylostella and can also inhibit the growth of Fusarium oxysporum at the concentrations higher than 0.39 mg/ml [13]. It has also been used for various pest controls, such as *Pieris rapae* larvae and *Plutella xylostella* [14].

Most of the *C. oleifera* shell and seed cake are always discarded or burned at present, causing serious environmental pollution and resource waste [15], and make it important to improve the utilization efficiency of *C. oleifera* waste and explore the methods of its utilization. Composting is one of the effective methods to treat *C. oleifera* shell and seed cake. However, the unstable components of the shell and seed, such as cellulose, hemicellulose and lignin, as well as saponins and tannins that can inhibit the growth of microbes and plant growth [3], significantly affect the biodegradation for their composting. Therefore, studying the degradation and content changes of tannin and saponin during the composting of *C. oleifera* shell and seed cake are necessary. In this study, the effects of the addition of *C. oleifera* seed cake on the composting of *C. oleifera* shell were investigated using the shell as the raw material and the seed cake as the microbial inoculants. The physicochemical properties of the compost and the degradations of tannin and saponin during the composting were analyzed, aiming to provide the scientific foundation for the composting of *C. oleifera* shell and seed cake.

## Materials and methods

### Experimental design

*C. oleifera* shell was collected from Zhejiang Jinhua Dongfanghong Forest Farm (China). The *C. oleifera* seed cake was provided by Zhejiang Tiantai Kangneng Tea Oil Co. Ltd. (China). The chemical compositions and primary properties of the raw materials are shown in Table 1 [3,16].

**Table 1.** Chemical compositions and properties of raw materials

| Parameters | *Shell of C.*<br>*oleifera* | Seed Cake of *C.*<br>*oleifera* | urea |
| --- | --- | --- | --- |
| Organic matter/% | 83.78 | 82.41 | - |
| Tannin/% | 2.26 | 1.03 | - |
| Saponin/% | 4.80 | 16.35 | - |
| Polysaccharids/% | 2.65 | 14.22 | - |
| Crude protein/% | 2.61 | 7.62 | - |
| Total C/% | 48.6 | 47.8 | 20 |
| Total N/% | 0.419 | 1.22 | 46.67 |
| C/N | 116 | 39 | 0.43 |
| Total P/% | 0.017 | 0.161 | 0 |
| **Total K/%** | 0.854 | 0.929 | 0 |
| **GI/%** | 58 | 0 | - |
| **pH** | 5.80 | 5.48 | 7.00 |

The uncrushed *C. oleifera* shell (partcle size≥20mm) and seed cake (particle size≤2mm) were composted in COMPOSTER 220^eco^ (73×115×80 cm, 220 L, BIOLAN), an insulated and highly ventilated ecological composting tank, in winter season with the ambient temperature varied from 0 to 22°C. Four treatments weere designed with the *C. oleifera* seed cake accounting for 1/3 (A1), 1/4 (A2), 1/5 (A3) and 1/10 (A4) of the total weight (dry weight), respectively, with three repetitions. The initial C/N ratio of each treatment was adjusted to 30 with urea and blended with 3% dry weight of EM bacteria. The water content of each treatment was then adjusted to 55%. The blends were stirred thoroughly and put in the composting tanks for aerobic fermentation. The compost temperature and room temperature were recorded every day at 3 pm for 76 days starting the first day of composting. Five hundred of grams compost were sampled on day 0, 7, 20, 30, 45, 60, and 90 by a multi-point sampling method from top, middle and bottom of the tank and measured for pH, the contents of tannin and saponin. The seed germination index (GI) assay was determined using seeds of *Brassica chinensis var* chinensis. All samples were kept in a −4°C refrigerator before the testing.

### Measurements of pH, conductivity and moisture content

Ten grams of fresh sample were dispersed in deionized water at the ratio of 10:1 (V (ml):W (g)) under ambient conditions. The suspension was shake at 200 r/min for 1 h and measured for pH with a pH −2F pH meter (Leici, Shanghai, China).

The moisture content was measured by drying at 105°C for 24h in a drying oven [3].

### Total organic carbon and elemental analysis

The total organic carbon (TOC) was determined using the protocol of “Physical and Chemical Analysis of Soils” (pg.376)[17]. The total nitrogen (N) using regular-kjeldahl method [18]; total phosphorus (P) and total potassium (K) were determined by spectrophotometry and flame photometry after H_2_SO_4_-H_2_O_2_ digest respectively [19].

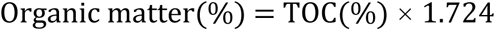

### Measurement of tannin and saponin contents

The contents of tannin was analyzed with spectrophotometry after extracted with a methanol-water mixture (1:1, v/v) [20]. And saponin were determined according to the methods [21].

### Seed germination index (GI)

Twenty grams of fresh sample were added into 200 ml of distilled water, thoroughly shaken for 1 h, leached at 30 °C for 24 h and filtered. Six milliliters of the filtrate were added into a 9 cm culture dish covered with a filter paper. Twenty good quality small green cabbage seeds were seeded in the culture dish and incubated in a 20 ± 1 °C incubator. The seed germination rate was measured in 24 h. Each treatment was repeated 3 times and distilled water was used as the control.

GI was calculated using the formula below.

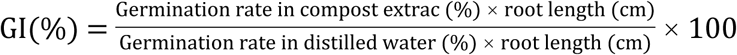

## Results and discussion

### Compost Temperature

Temperature is one of the important compost maturity indexes. It is closely related to the properties of raw material, microbial activity, composting cycle and compost maturity [22,23]. In general, a composting undergoes four phases including initial mesophilic phase, thermophilic phase (>50°C), cooling phase and maturing phase. Temperatures over 55°C kill pathogenic microorganisms, eggs and weed seeds, producing harmless composts. However, microbial activities drops rapidly at the temperatures above 63°C [24]. The initial temperatures for the four treatments were all measured to be 10 ± 0.8°C (Fig 1). The temperature reached over 50°C on day 2 (treatment A1) or day 3 (the other treatments). The numbers of the days with the temperatures higher than 50°C were 24, 20, 17 and 17, respectively for the treatments A1, A2, A3 and A4 during the 90 day’s composting, with the highest temperature records of 71.5°C, 66°C, 63°C and 62°C. The numbers of the days with the temperatures higher than 55°C were 20, 18, 10 and 3 for the treatments A1, A2, A3 and A4, respectively. The longest duration with the temperature above 55°C were 11 days, 7 days, 6 days and 3 day for the treatments A1, A2, A3 and A4, respectively. These observations suggest that the content of *C. oleifera* seed cake significantly affects the compost heating. The higher contents of seed cake in the compost cause faster temperature rising, longer thermophilic phase and higher maximum temperatures, and vice versa. As shown in Table 1, the seed cake contains much more polysaccharides and crude protein than the shell. The contents of polysaccharides and crude protein in the four treatments are in the order of A1>A2>A3>A4. Therefore, the duration of thermophilic phase and the highest temperature are probablely affected by the contents of polysaccharides and crude protein in *C. oleifera* shell and seed cake.

**Fig 1.**
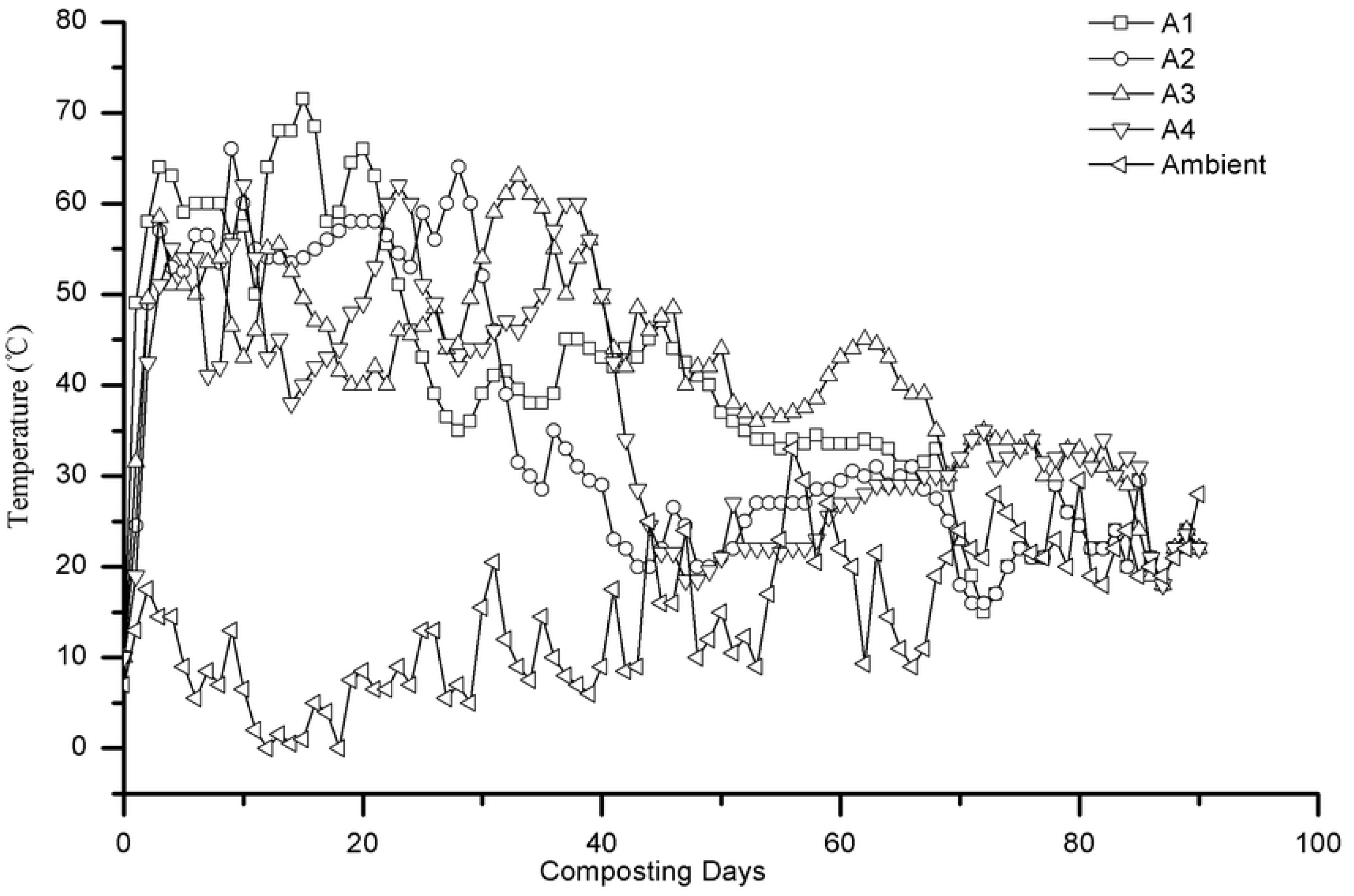
Temperature profiles of the compost during composting.

### pH

pH is one of the important factors affecting the growth and reproduction of microorganisms. The optimum pH for the growth of most microorganisms is 6.5-7.5. The pHs of treatments A1, A2, A3 and A4 respectively increased from the initial values of 5.82, 5.98, 6.04 and 6.06 to 7.86, 7.73, 7.92 and 7.74 in day 30, and slightly decreased to the values of 7.65, 7.60, 7.22, and 7.06 on day 60, and remained stable thereafter (Fig 2). Due to the shell and seed cake of *C. oleifera* in faintly acid (Table 1), the initial pHs of the composts are acidulous. At the early composting stage, the microbes rapidly decomposes the organic nitrogen in the compost, resulting in a large amount of ammonia, and leading to alkaline pH values [25]. With the increase of temperature, the ammonia is continuously volatilized and simultaneously, organic acids are continually produced, leading to the decrease of pH value in the composts [26]. At the cooling stage of compost, the organic acids are gradually decomposed or converted into complex cyclic compounds such as humus, which moves the composting to mature, and the stack pH becomes stable consequently [24].

**Fig 2.**
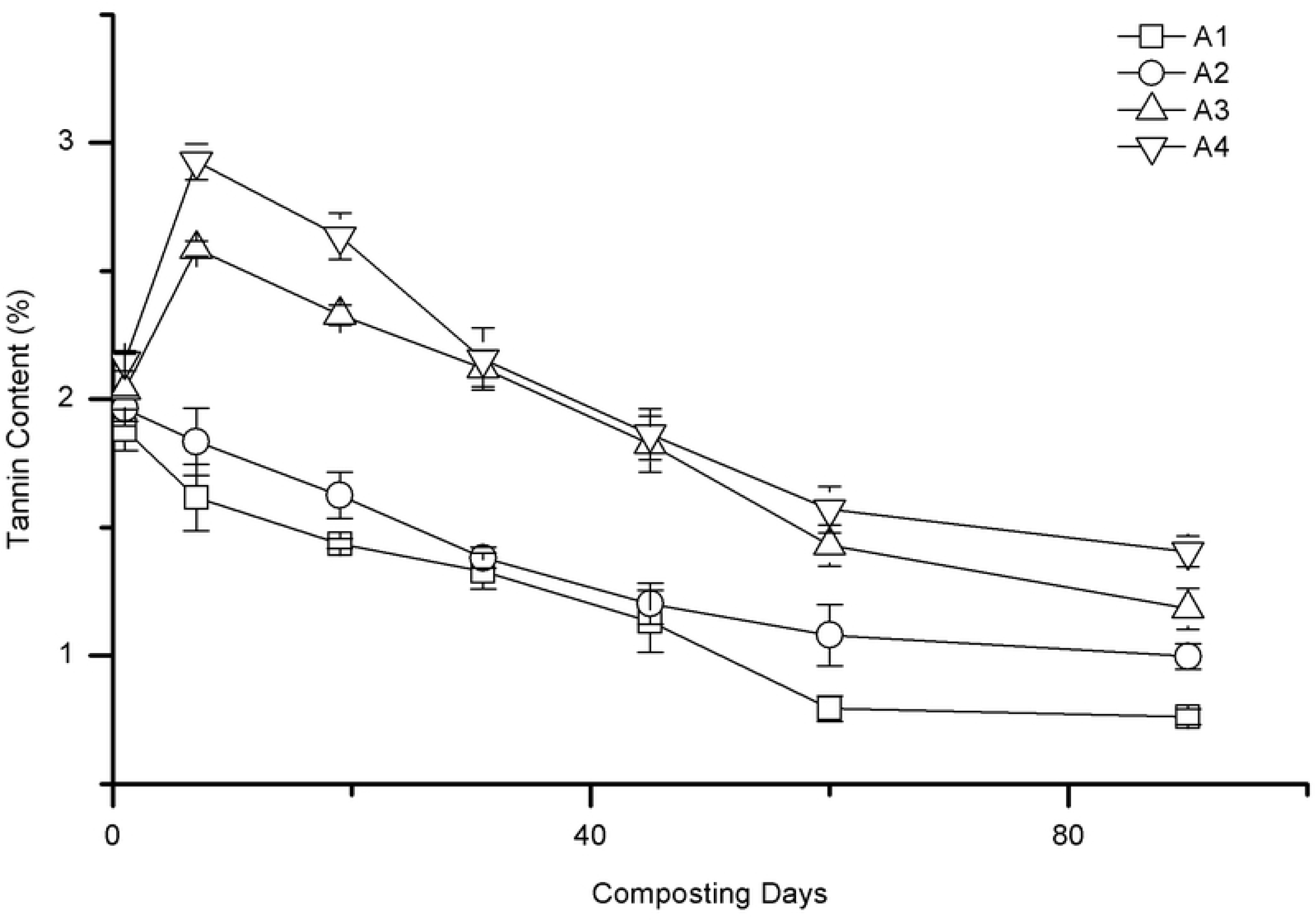
Changes of pH during composting.

### Tannin and Saponin

The tannin contents of treatments A1 and A2 exhibited similar trend that both of them decreased linearly with the composting time (Fig 3). Those of A3 and A4, however, presented another trend that they increased in the early 7 days and declined rapidly thereafter. At the end of the experiment, the tannin contents of the four treatments are in the order of A1<A2<A3<A4 with the values of 0.761%, 0.997%, 1.183% and 1.406%, respectively (Table 1), and the corresponding degradation rates of 59.52%, 49.20%, 41.89%, and 34.57% (Fig 4). As the proportion of the seed cake increased in composting, the tannin content in the final compost product decreased, and the tannin degradation rate increased, which were affected significantly (P<0.05) by the the proportion of the seed cake and might be attributed to combination of protein and tannin. Protein is an important nutrient for microorganisms to provide heat for the compost to enter thermophilic phase [24]. Tannin can combine with protein to form insoluble complex, which affects the degradation ability of microorganisms and the degradability of protein in the compost [27]. At the beginning of this experiment, the crude protein contents of the four treatments are in the order of A1>A2>A3>A4, and their tannin contents follow the order of A4>A3>A2>A1 (Table 1). Therefore, the treatments with higher tannin contents has longer initial phase and shorter duration of thermophilic phase.

**Fig 3.**
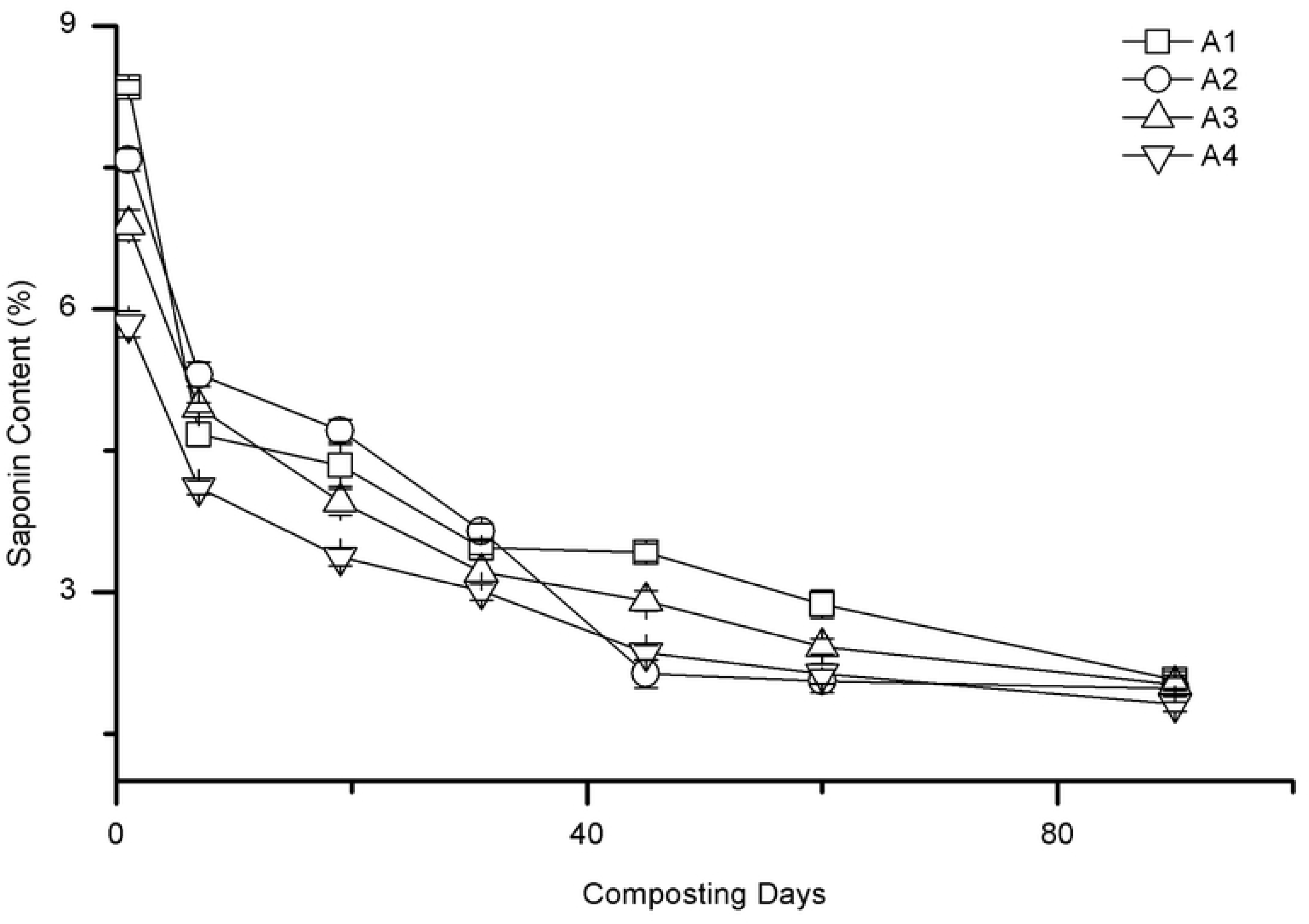
Changes of Tannin content during composting.

**Fig 4.**
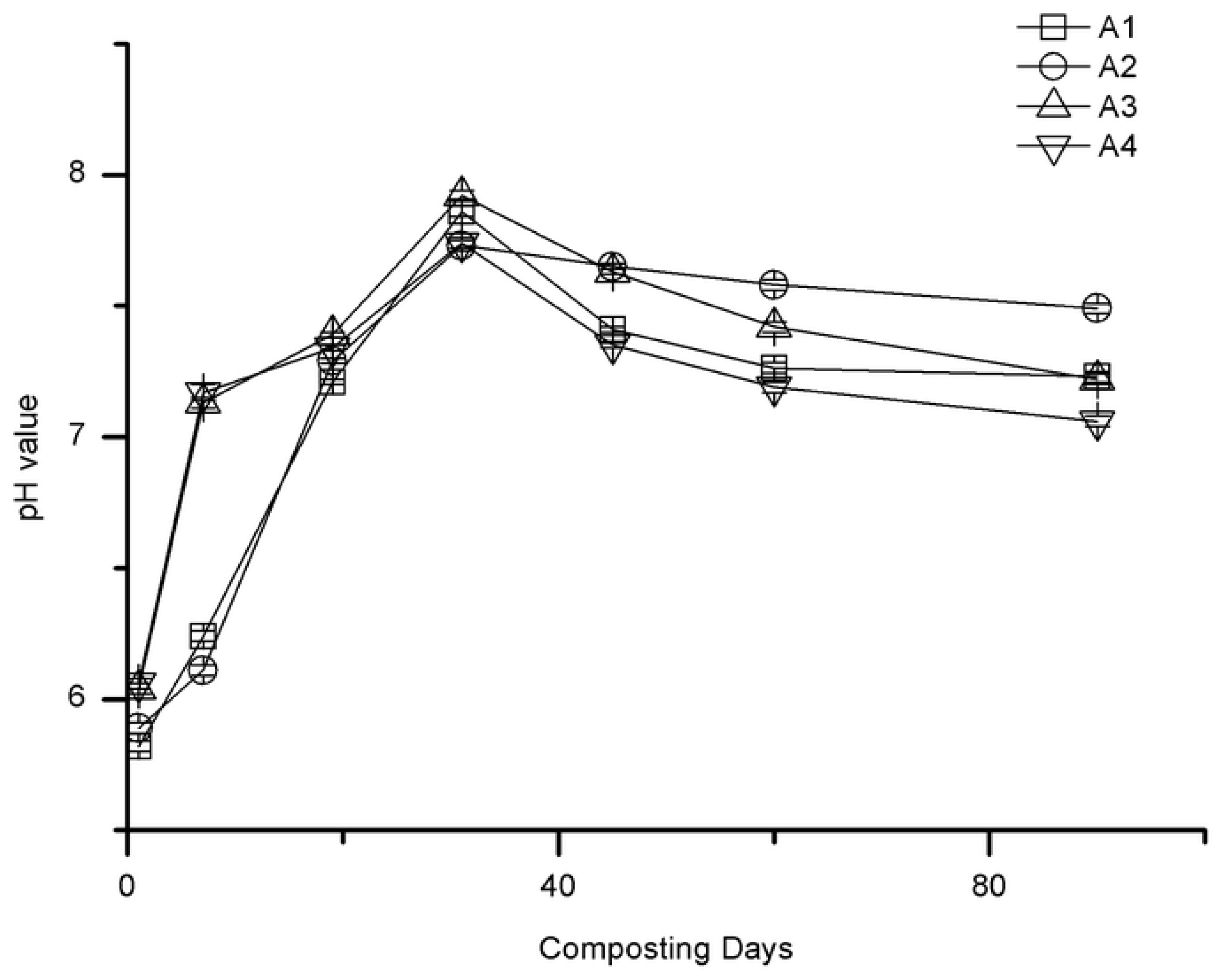
Degradation rates of tannin and saponin during composting.

Fig 5 shows the changes of the saponin contents in the four experiment groups with composting time. The initial saponin contents are in the order of A1>A2>A3>A4, and exhibited decreasing trends with the composting time. The saponin degradation rate increased with the increase of seed cake content. The final degradation rates of the group A1, A2, A3 and A4 were determined to be 75.22%, 73.92%, 70.65%, and 68.92%, respectively. With the increased content of the seed cake, the saponin degradation rate of the four treatments decreased, but not the differences were not obviously. The final saponin contents in the compost products were all about 1.81-2.07%, and no correlation with the initial addition of *C, oleifera* seed cake.

**Fig 5.**
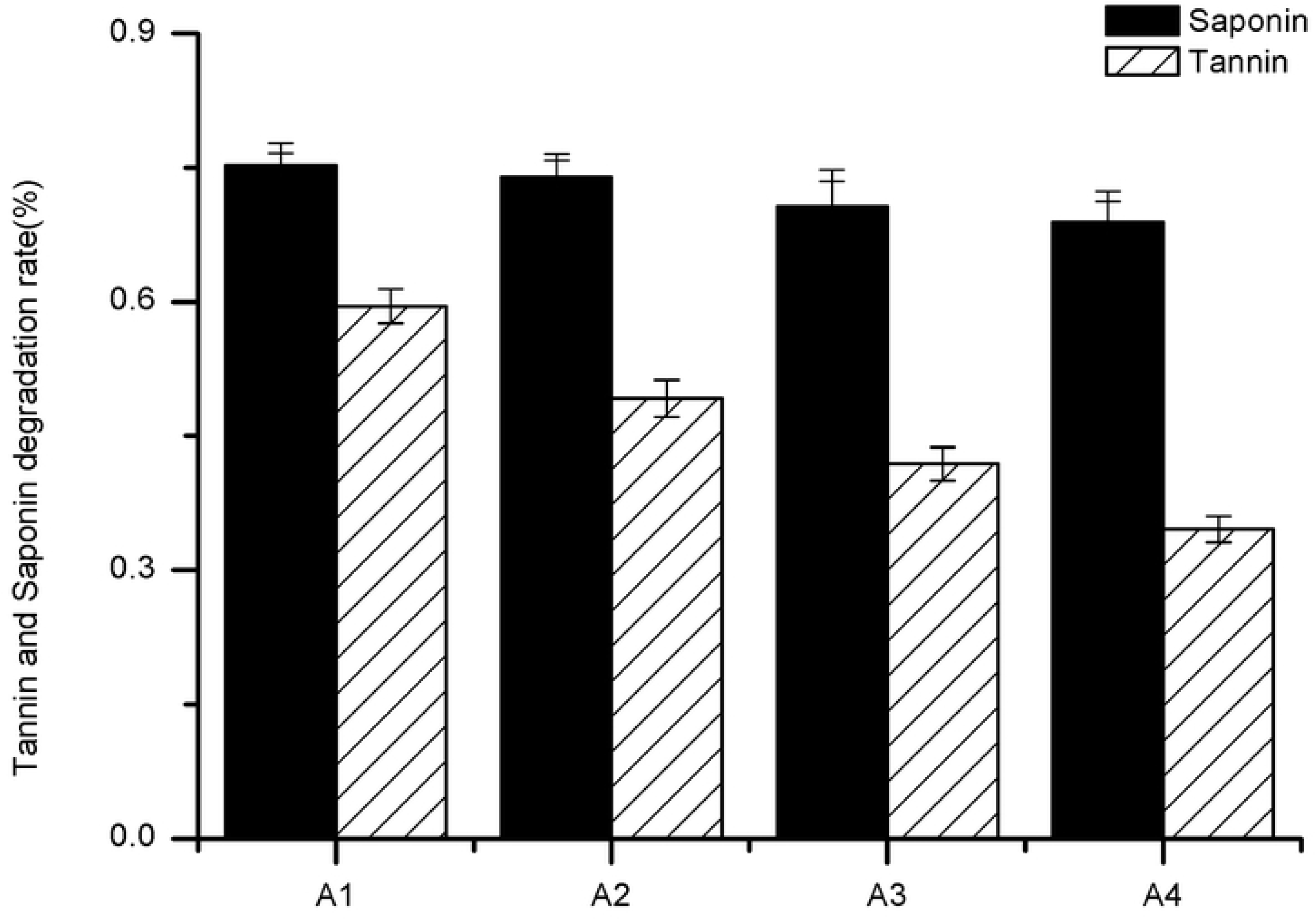
Changes of saponin content during composting.

### Chemical composition

Table 2 lists the chemical properties of the final compost product of each group in 90 days. The moisture contents in the compost products of all four groups are less than 30%. The organic matter contents are greater than 50% and the pH are between 7.06 and 7.61. The GI values are greater than 122%, which means the final compost product is safe for the growth of plants. The total nutrient contents (N, P_2_O_5_ and K_2_O) of the four groups are less than 5%, increased by 2.67, 2.06, 2.03, 1.95 times as much as the initial total nutrient contents of A1, A2, A3 and A4, respectively. The contents of organic carbon and nitrogen of all four groups were decreased by the compost because carbon was consumed and converted into carbon dioxide and humus and nitrogen was volatilized as ammonia, or converted into nitrate and nitrite, or assimilated by organisms [28,29]. Compost maturity takes place at C/N<20, and the higher the degree of maturity, the closer the C/N ratio is to 10 (the C/N ratio of humus) [30]. The C/N ratios of the compost products of A1, A2, A3 and A4 are 18.52, 19.71, 21.55 and 30.52, respectively. The product of A1 met the standard of compost maturity most.

**Table. 2.** Chemical properties of the composts

| The initial chemical properties of the compost |  |  |  |  |  |
| --- | --- | --- | --- | --- | --- |
| Parameters | A1 | A2 | A3 | A4 |  |
| Moisture(%) | 55 | 55 | 55 | 55 |  |
| Organic matter (%) | 83.44 | 83.51 | 83.55 | 83.66 |  |
| Total N(%) | 0.619 | 0.579 | 0.553 | 0.492 |  |
| Total P<sub>2</sub>O<sub>5</sub>(%) | 0.121 | 0.105 | 0.094 | 0.069 |  |
| Total K<sub>2</sub>O(%) | 1.052 | 1.047 | 1.044 | 1.037 |  |
| Total nutrients(%) | 1.79 | 1.73 | 1.69 | 1.59 |  |
| pH | 5.82 | 5.89 | 6.04 | 6.06 |  |
| C/N ratio | 30 | 30 | 30 | 30 |  |
| GI(%) |  |  |  |  |  |
| The final chemical properties of the compost |  |  |  |  | China<br>organic<br>fertilizer |
| Parameters | A1 | A2 | A3 | A4 |  |
| Moisture(%) | 26.5±1.1 | 25.8±0.9 | 25.3±1.2 | 25.1±0.4 | ≤30 |
| Organic matter<br>(%) | 51.36±2.1 | 52.13±1.8 | 50.72±2.2 | 50.44±1.9 | ≥45 |
| Total N(%) | 2.51±0.06 | 1.64±0.04 | 1.84±0.05 | 1.46±0.05 | - |
| **Total P<sub>2</sub>O<sub>5</sub>(%)** | 0.34±0.04 | 0.25±0.03 | 0.17±0.02 | 0.12±0.03 | - |
| **Total K<sub>2</sub>O(%)** | 1.95±0.05 | 1.68±0.06 | 1.43±0.04 | 1.52±0.04 | - |
| **Total nutrients(%)** | 4.79 | 3.57 | 3.44 | 3.10 | ≥5 |
| **pH** | 7.23±0.04 | 7.61±0.05 | 7.22±0.03 | 7.06±0.07 | 5.5-8.5 |
| **C/N ratio** | 18.52 | 19.71 | 21.55 | 30.52 | - |
| **GI(%)** | 124.03±2. | 137.53±4. | 158.72±2. | 122.31±2. | ≥85 |
|  | 5 | 3 | 6 | 2 |  |

These results indicate that the nutrient content is positively correlated with the initial seed cake content and negatively with the C/N ratio of the final compost product (Tablae 3). The higher the content of the seed cake lead to the higher degree the compost maturity is. Overall, the compost products meet the criteria of the Chinese Standard of Organic Fertilizer NY 525-2012.

## Conclusion

The duration of thermophilic phase and highest temperature during the co-composting are mainly affected by the addition of seed cake for its content of polysaccharides and crude protein. The addition of seed cake is positively correlated with the degradation ratio of tannin and saponin and negatively correlated with the final tannin content and C/N ratio. The compost obtained was 50.44-52.13% of organic matter, 3.10-4.79% of total nutrients (N, P_2_O_5_ and K_2_O), 1.81-2.07% of saponin, 7.06-7.61 of pH and 122.31-158.72% of GI. Overall, the addition of seed cake promoted the stability, fertilizer efficiency and safety of the compost product.

## Acknowledgements

This work was supported by the Provincial Department of Science and Technology of Zhejiang, China, Grant No. 2017C02022.

